# Social and physical environment independently affect oviposition decisions in *Drosophila melanogaster*

**DOI:** 10.1101/2021.01.27.428449

**Authors:** Emily R. Churchill, Calvin Dytham, Jon R. Bridle, Michael D.F. Thom

**Affiliations:** School of Biological and Marine Sciences, University of Plymouth, Plymouth UK; Department of Biology, University of York, York UK; Department for Genetics, Evolution and Environment, University College London, London UK

## Abstract

In response to environmental stimuli, including variation in the presence of conspecifics, animals show highly plastic responses in behavioural and physiological traits influencing reproduction. These responses have been extensively documented in males, but equivalent study of females is so far lacking. We expect females to be highly responsive to environmental variation, with significant impacts on fitness given females’ direct impact on offspring number, size, and developmental conditions. Using *Drosophila melanogaster* as a model, we manipulate (a) exposure to conspecific females, expected to influence their expectation of number of potential mates and larval density for their own offspring, and (b) test how prior consexual population density interacts with the spatial distribution of potential oviposition sites, with females expected to prefer clustered food resources that can support a larger number of eggs and larvae. After exposure to competition, females were slower to start copulating and reduced their copulation duration – the opposite effect to that observed in males previously exposed to rivals. There was a parallel and perhaps related effect on egg production, with females previously housed in groups laying fewer eggs than those that were housed in solitude. The spatial distribution of resources also influenced oviposition behaviour: females clearly preferred aggregated patches of substrate, being more likely to lay, and laying on more of the available patches, in the clustered environment. However, we found no significant interaction between prior housing conditions and resource patchiness, indicating that females did not perceive the value of different resource distributions differently when they were expecting either high or low levels of larval competition. While exposure to consexual competition influences copulatory behaviours, it is the distribution of oviposition resources that has a greater impact on oviposition decisions.

## Introduction

### Effects of intra-sexual competition

Most individuals are likely to experience competition for resources during at least some part of their lifetime. While population density is partly responsible for determining the extent of competition, the distribution of resources in the environment also plays an important role: more clustered resources (whether food, mates, or nesting/oviposition sites) result in increased encounter rates (Emlen and Oring, 1977) and thus a greater degree of competition. Because optimal responses often differ in high- and low-competition environments (Maynard Smith, 1979), animals which experience variation in local population density are expected to make plastic adjustments to behaviour and physiology in response to prevailing levels of competition, in order to maximise their lifetime reproductive success. The effect of exposure to conspecific rivals on reproductive investment has been particularly extensively studied in this regard, most extensively among males (Klemme and Firman, 2013, Rowley et al., 2019, Droge-Young et al., 2016, Gage and Barnard, 1996, García-González and Gomendio, 2004, Gomendio and Roldan, 1991, Kiss et al., 2019, Kustra et al., 2019). *Drosophila* species have emerged as an important model organism for understanding these effects: males of these species are sensitive to the presence of potential competitors and adjust a range of behaviours and reproductive physiological process accordingly (Bretman et al., 2009, Fedorka et al., 2011, Garbaczewska et al., 2013, Moatt et al., 2014).

Surprisingly, far less research attention has focused on the equivalent plasticity in females (but see Singh and Singh (2014) on density effects on female remating rates, and Battesti et al. (2012) and Sarin and Dukas (2009) on oviposition copying behaviours). Female *Drosophila* are aggressive towards other females, initiating a range of behaviours similar to those observed among fighting males (Bath et al., 2017, Chapman and Wolfner, 2017, Ueda and Kidokoro, 2002), suggesting that females may be equally as sensitive to the presence of same-sex rivals as are males. However, while female intrasexual aggression is common, they also show strong social attraction to conspecifics on food patches (Lihoreau et al., 2016), perhaps because of the efficiency benefits of shared feeding among larvae (Dombrovski et al., 2017). This attraction remains to be fully explained, as there is inevitably a trade-off between these benefits and increasing competition, which can even lead to cannibalism (Vijendravarma et al., 2013). This tension between competition and cooperative feeding is to some degree mediated by relatedness, with closely related larvae more likely to form cooperative feeding aggregations than are unrelated larvae (Khodaei and Long, 2020). Nevertheless, under food restriction cannibalism is observed even within inbred laboratory strains with high mean relatedness (Vijendravarma et al., 2013).

### Plasticity in oviposition decisions

During oviposition females can only assess the level of competition their larvae will face based on the number of existing eggs at a patch, or the number of pheromone markings by conspecifics (Malek and Long, 2020, Tait et al., 2020). It seems likely that they may also be sensitive to intra-sexual encounter rate among adults as a proxy for likely larval competition environment. However despite the strong evidence for density and encounter rate effects on male behaviour (Churchill et al., 2020, Bretman et al., 2009, Moatt et al., 2014, Fedorka et al., 2011, Price et al., 2012, Hopkins et al., 2019, Garbaczewska et al., 2013), and evidence that females are sensitive to the presence of conspecifics when making oviposition decisions (Malek and Long, 2020, Tait et al., 2020), studies on the effect of female encounter rate on subsequent behaviour are so far lacking (Gillmeister, 1999).

Here we test whether females respond to the presence of other females during adulthood by subsequently plastically adjusting their egg laying behaviour according to expected level of larval competition. Wild female *D. melanogaster* lay on rotting and fermenting fruits, a naturally patchy and ephemeral environment. Females make sophisticated egg laying decisions, including assessing not only the nutritional quality of the resource but also the inter-patch substrate, considering potential energetic costs of larval travel (Schwartz et al., 2012). As they lay only one egg at a time (Yang et al., 2008), evidence of clustered eggs shows evidence of repeated decisions to lay in the same site. As well as manipulating adult density, we tested how any socially-induced plasticity interacts with the physical oviposition environment. We hypothesized that females would lay fewer eggs per patch on small, isolated patches, and more eggs per patch on clustered patches as this better facilitates larval travel between food sources. Furthermore, we expected that this would be mediated by the female’s experience of intra-sexual encounter rates prior to egg laying, with females from a high-density environment laying investing more resources per egg, and thus laying fewer eggs overall, in the expectation of high future competition among larvae.

## Methods

All fly rearing and experiments were conducted at 25°C on a 12hour light:dark cycle (08:00 – 20:00h GMT), unless otherwise stated. Stock flies originating from a Canton-S laboratory stock population were housed in 40ml vials containing 7ml of a standard agar-based medium (40g of yeast and 40g sucrose per litre); hereon in described as standard vials. Approximately 25 *D. melanogaster* were held in each vial, and all vials were pooled and randomly redistributed into new vials every seven days to minimise any within-vial effects of inbreeding, drift, and selective sweeps.

Parental generation vials were set up with a standardised density of six males and six females per vial to ensure food resources were not limiting. Test flies were offspring collected under ice anaesthesia from these parent vials within six hours of eclosion to ensure virginity, and immediately transferred into treatment conditions.

### Prior experience of competition

Test females were housed in treatment vials for seven days in one of two treatments: singly housed (hereafter “solitary”), or in a group of six females (“grouped”).

### Copulation behaviours

Females were translocated to a new standard vial for copulation with a standard seven-day old male which had previously been housed alone since eclosion. Courtship and copulation behaviours were observed live, and latency to copulate (time from when pair were first introduced until the male successfully mounts the female) and copulation duration (until pair fully separates) were recorded in seconds.

### Oviposition distributions

Females that did not copulate within 90 minutes of being introduced to the mating vial were excluded from subsequent (egg laying) stages of the experiment. Females which copulated were transferred to individual egg laying dishes, with oviposition substrate arranged in one of two spatial treatments: dispersed or clustered resources. Petri dishes were 140mm in diameter and contained four patches of agar-based medium (each 22mm in diameter, 7mm depth). For the dispersed treatment, patches were located at four equidistant points around the circumference of the petri dish, at an interpatch distance of 100mm (Fig. 1a). In the clustered treatment, patches were arranged in a square in the centre of the Petri dish (Fig. 1b). Each patch was placed 3mm from the edge of the Petri dish, or from other patches, to keep total surface area available for oviposition constant between treatments. The base of each Petri dish was lined with filter paper, to which 10ml of distilled water was added to prevent food patches from drying out.

**Figure 1.**
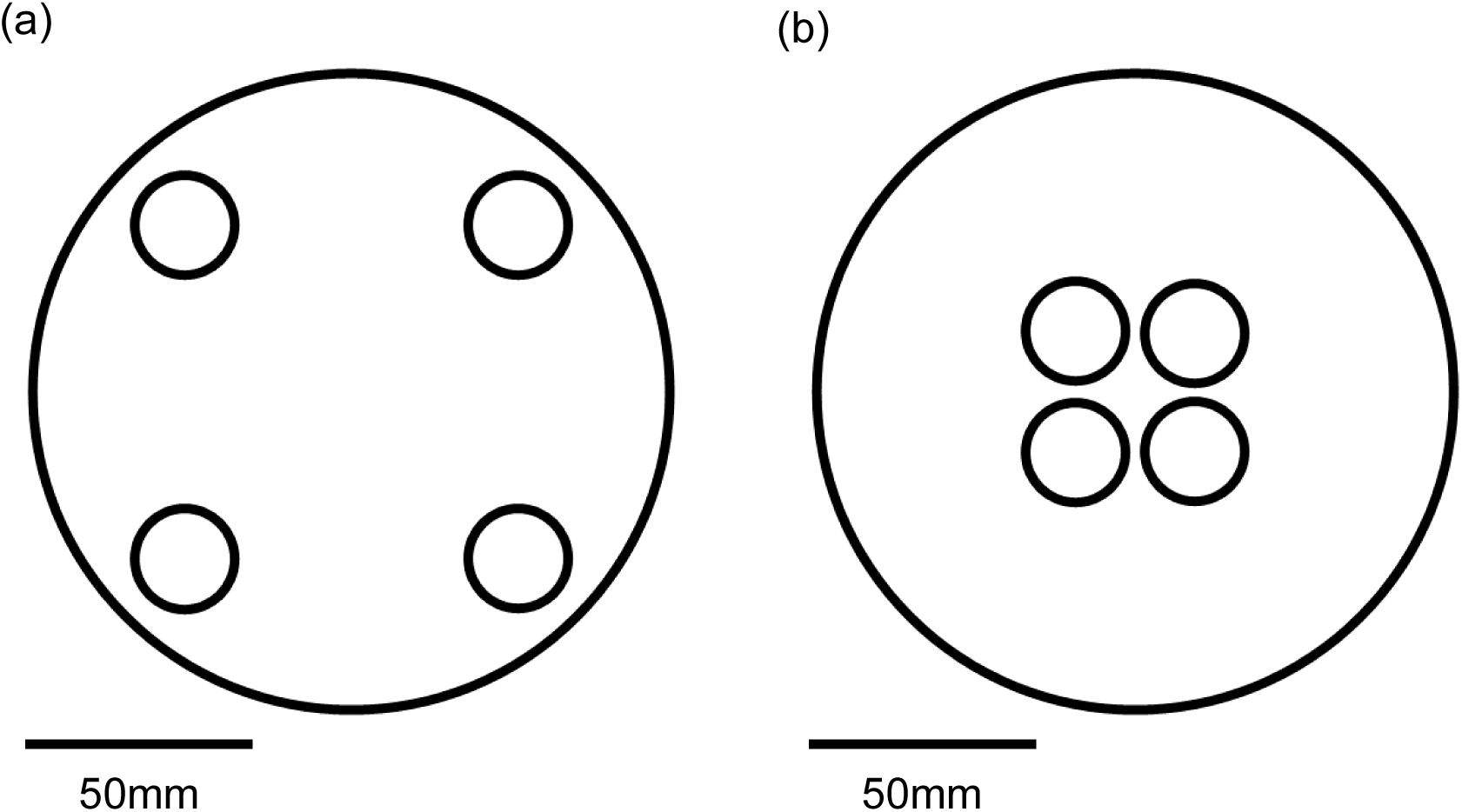
(a) Dispersed resource distribution treatment: food discs located at four equidistant points around the circumference of the Petri dish. (b) Clustered resource distribution treatment: food discs located in the centre of the Petri dish, in a square arrangement with each disc approximately 3mm apart from adjacent discs.

This arrangement resulted in four treatments: females from solitary and grouped treatments could lay in either dispersed or clustered resource plates in a fully factorial design. Sample sizes were: solitary/clustered resources: 22; solitary/dispersed resources: 22; grouped/clustered resources: 30; grouped/dispersed resources: 29. For the analysis of mating behaviours only, we had an additional six solitary females for which there is no accompanying egg laying data due to incubator failure.

Females were left to oviposit eggs in these dishes for 18-20 hours. Treatment enclosures were placed randomly in three incubators, maintained at 25°C, under constant light to allow imaging. Each incubator held one Raspberry Pi (www.raspberrypi.org) connected to an 8MP Raspberry Pi Camera module (v2; www.thepihut.com). Frame capture software ‘raspistill’ was used to capture one image every 10 minutes. For each image, we recorded on which patch of the four food patches the female was found, or if she was not currently on a patch.

Once the female had been removed, egg-laying plates were photographed using a digital camera (Panasonic Lumix DMC-FT4) to allow counting of eggs laid per patch.

### Fitness

The plates containing eggs were returned to the incubator, and after 21 days the emerged adults were counted and sexed. During the 21 day emergence period, the filter paper was replenished with 5ml of water every three to four days.

Immediately after females were removed from the plates (18-20h after introduction), they were given a further seven days to lay any remaining eggs in a standard vial in order to quantify, before being removed for wing size measurements to be taken. Number of male and female offspring were counted 14 days later.

### Statistical analysis

All statistical analyses were conducted in R v4.0.3 (R Core Team, 2019).

We tested the effects of treatment on response variables using mixed effects models with the appropriate error distribution (binomial error for egg presence/absence, negative binomial error distribution for the overdispersed egg and offspring number, Gaussian for mating latency and duration) with the functions in packages lmerTest (Kuznetsova et al., 2017), and lme4 (Bates et al., 2015). This approach allowed us to fit vial identity as a random effect to account for shared housing of females in grouped treatment. However, in all models, the variance component for vial was estimated at non-significantly different from zero leading to a singular model fit, so we re-ran these using (generalized) linear models with the appropriate error distribution: binomial for the egg presence/absence data, quasi-Poisson error for the egg and offspring number models to account for overdispersion, and Gaussian for mating latency and duration.

We analysed whether the number of patches a female chose to lay on was influenced by either prior housing or oviposition substrate treatment using Chi-squared tests. We recognize that this does not allow us to treat the effect of shared housing as a random effect, although this was not a significantly confounding factor in any of the previous models.

## Results

### The impact of prior exposure to density on female mating behaviours

All females were courted; all group-housed females and all but 3 out of 60 single-housed females copulated, indicating that prior housing had no effect on whether females received courtship or subsequently copulated.

Group-housed females were >120s slower to copulate than solitary females (latency to mate, solitary: 235 ± 47s; group housed: 373 ± 45s; linear model: log_10_(copulation latency): F_1,104_ = 15.97, p < 0.001; Fig. 2). The residuals from this model were non-normal (Shapiro-Wilk test p = 0.010) due to outliers in both treatments (Fig 2). Removing all values > 1000s (N = 4) normalized the residuals (p = 0.820) and the difference between treatments remained statistically significant (F_1,100_ = 17.61, p < 0.0001).

**Figure 2.**
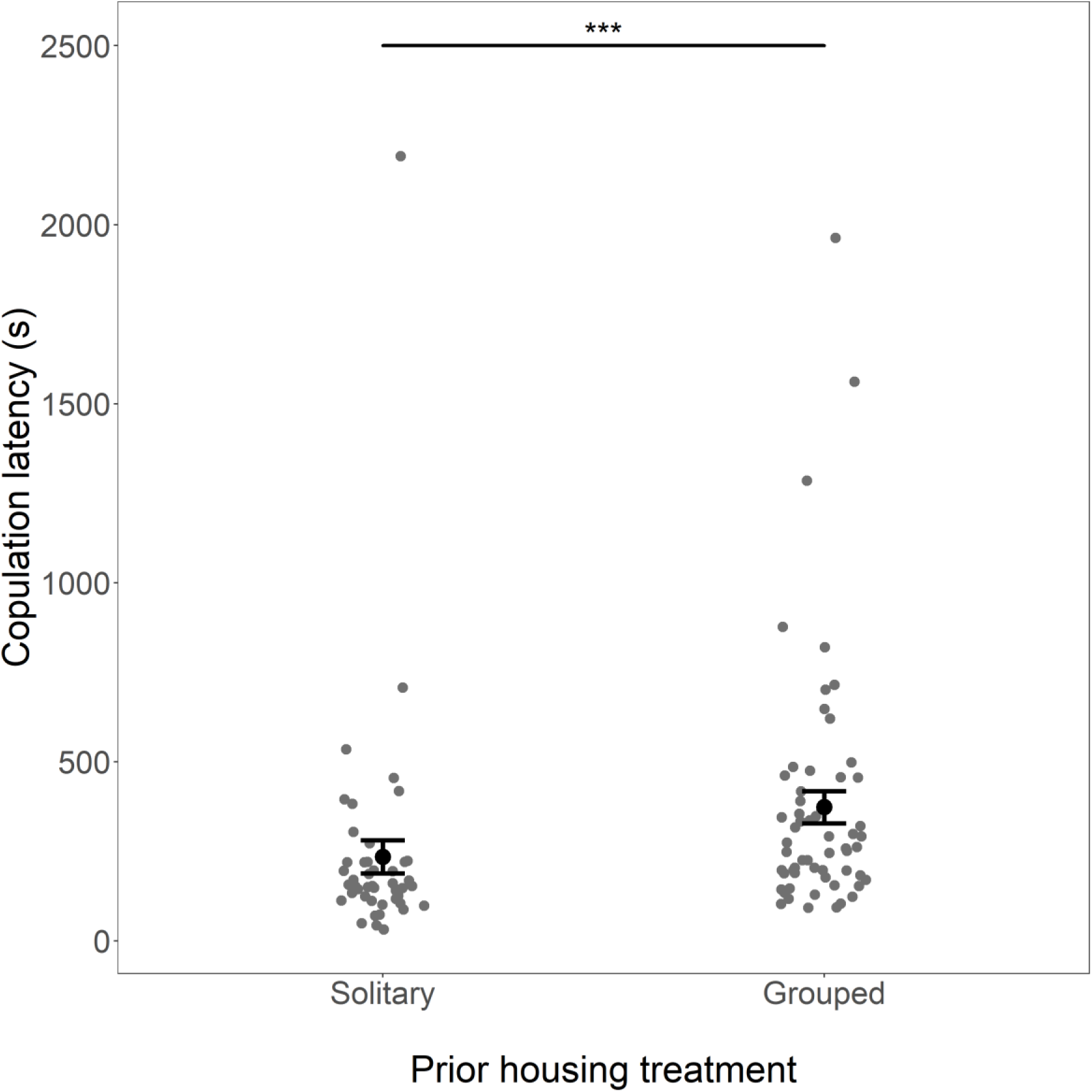
The effect of female prior housing density on latency to copulate, shown as means (black dot) and 95% confidence interval of copulation latency in seconds. The difference in latency remained statistically significant after removal of the four points with values above 1000s. Note that the analysis was performed on logged data, but untransformed values are presented here.

As well as being slower to start copulating, co-housed females’ mating duration was an average 55 seconds less than that of solitary females (linear model: log_10_(mating duration): F_1,102_ = 4.97, p = 0.0255; Fig. 3). This difference remained significant after removal of an outlier in the group-housed group (linear model; F_1,101_ = 4.10, p = 0.0418; Fig. 3).

**Figure 3.**
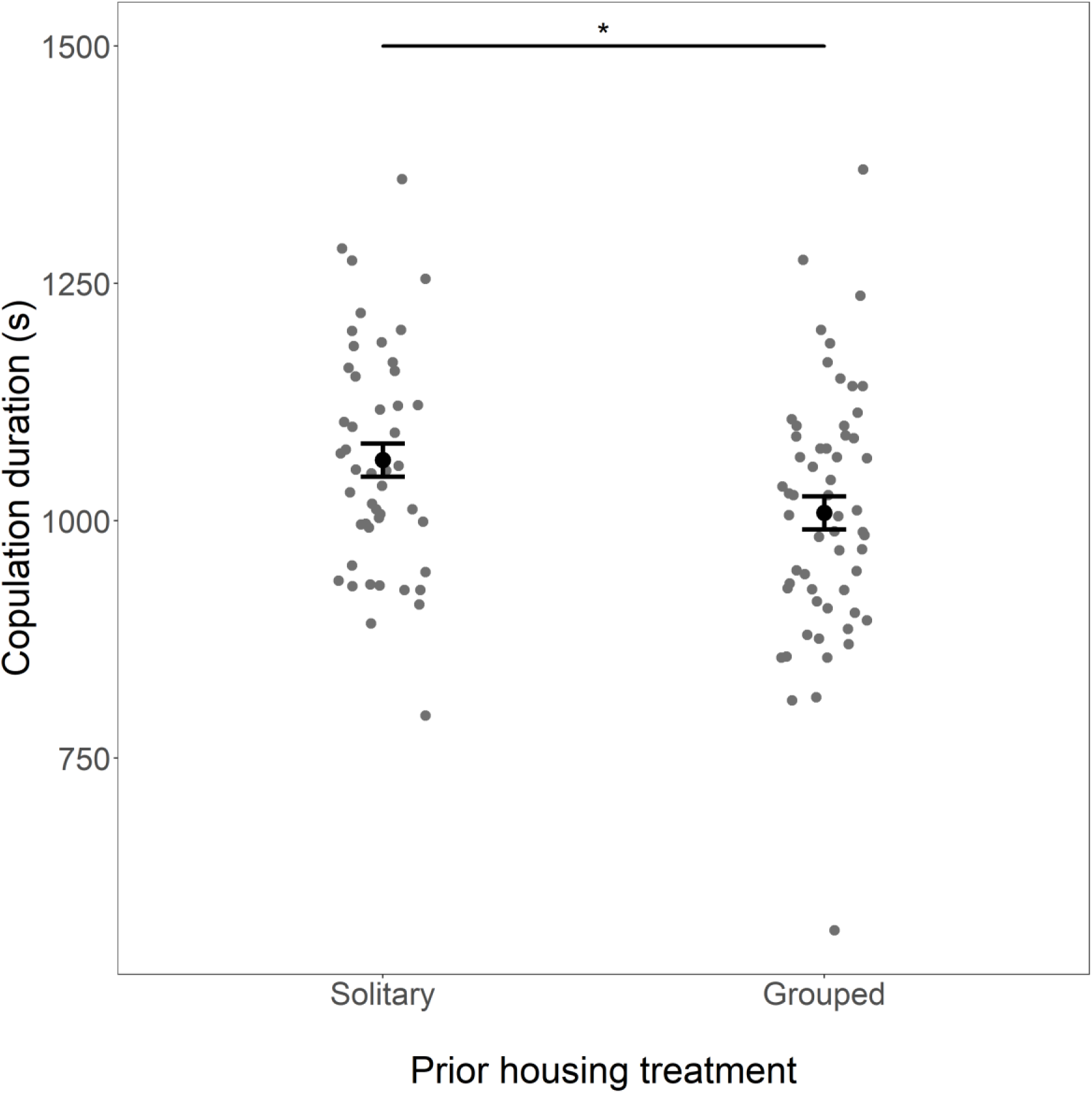
The effect of female prior exposure to competition on copulation duration. Means (black dot) and 95% confidence interval of copulation duration are shown in seconds. The difference in means remains significant after removal of the low outlier in the group-housed treatment. Note that the analysis was performed on logged data, but untransformed values are presented here.

**Figure 4.**
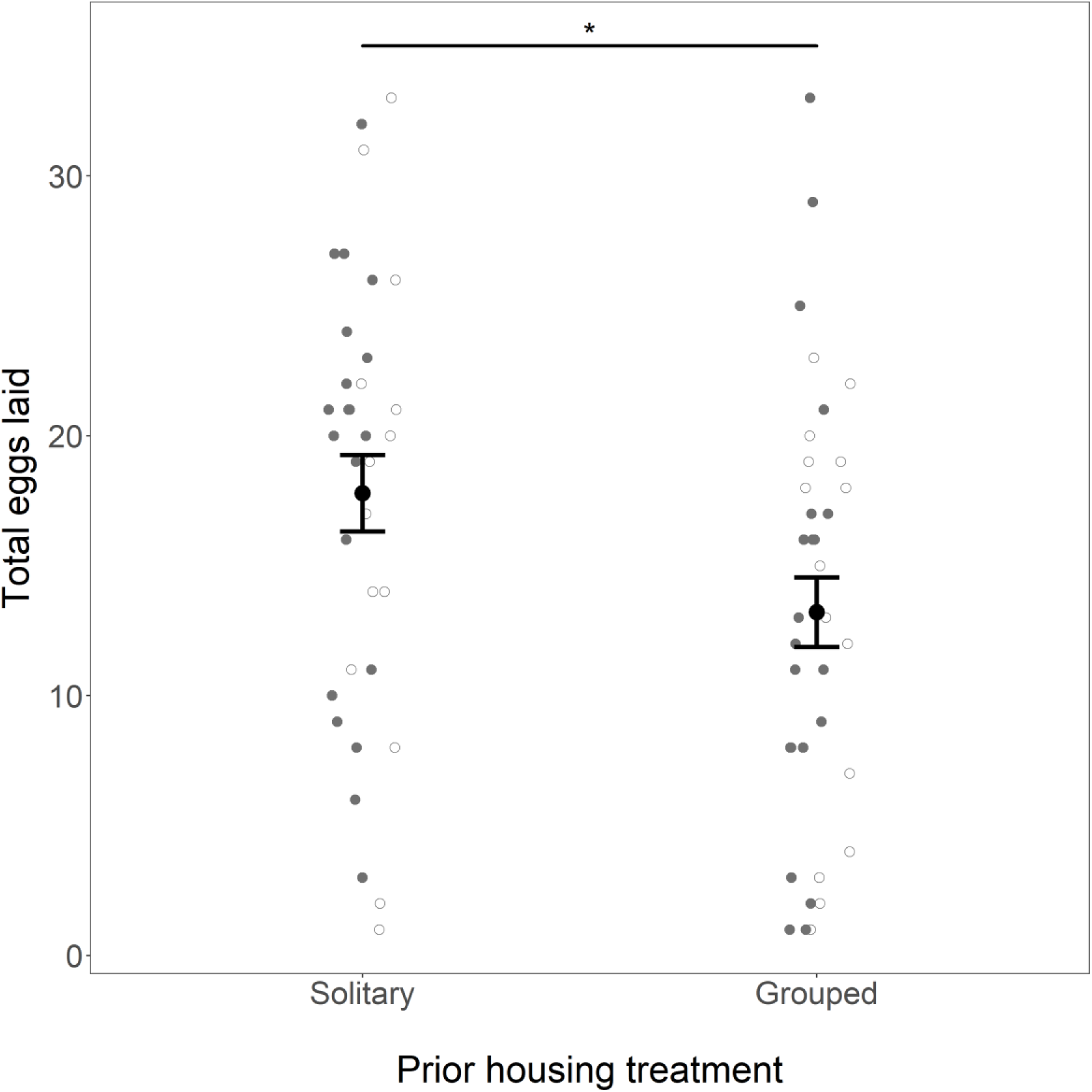
Prior housing density significantly influenced how many eggs were laid, with females from a solitary background laying more eggs than those that had previously been group housed. Resource distribution, by comparison, had no significant effect on laying rate (filled points: clustered resources; open points: dispersed resources). Means (black dot) and 95% confidence interval are shown for the two housing densities.

**Figure 5.**
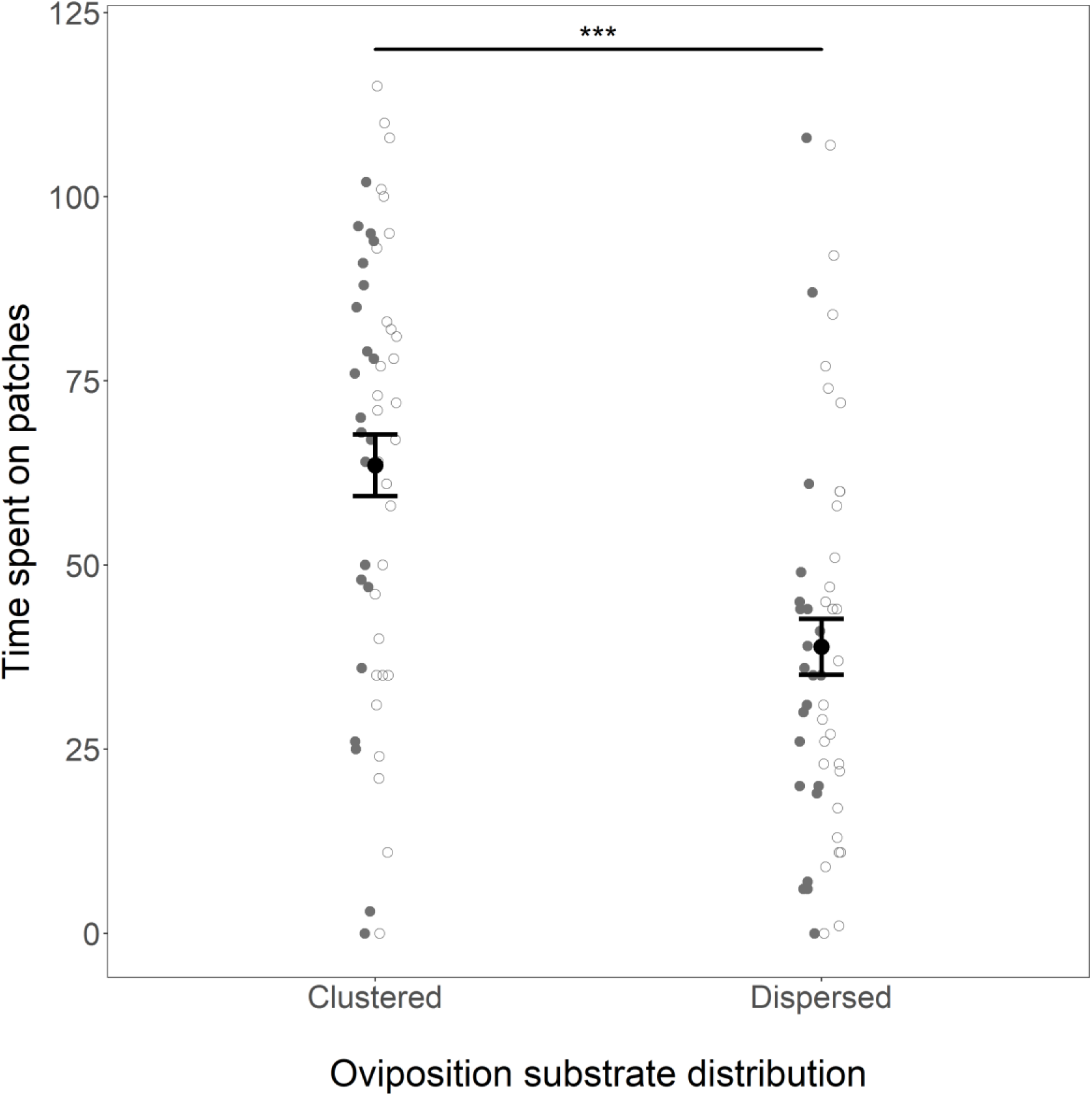
The effect of the spatial distribution of oviposition resources on the time females spent visiting those resources (measured as the number of images in which the female was observed on a food patch). Means (black dot) and 95% confidence intervals are shown. Filled points – grouped females; open points – solitary females.

### Effects of prior housing density and resource spatial distribution on oviposition and fitness

Female oviposition behaviour could be influenced by two treatment conditions: prior housing (either group-housed or solitary) and current oviposition landscape (either clustered or dispersed food patches). Of these, clustered resources led to a significantly higher proportion of females laying than did dispersed resources (general linear model with binomial errors; X^2^ = 4.88, p = 0.0272; Table 1), but no significant interaction between these variables influenced whether eggs were laid or not (X^2^ = 1.24, p = 0.265), and while the percentage of females laying eggs was depressed among group-housed females, prior housing treatment had no significant effect (X^2^ = 2.74, p = 0.098; Table 1).

**Table 1.**
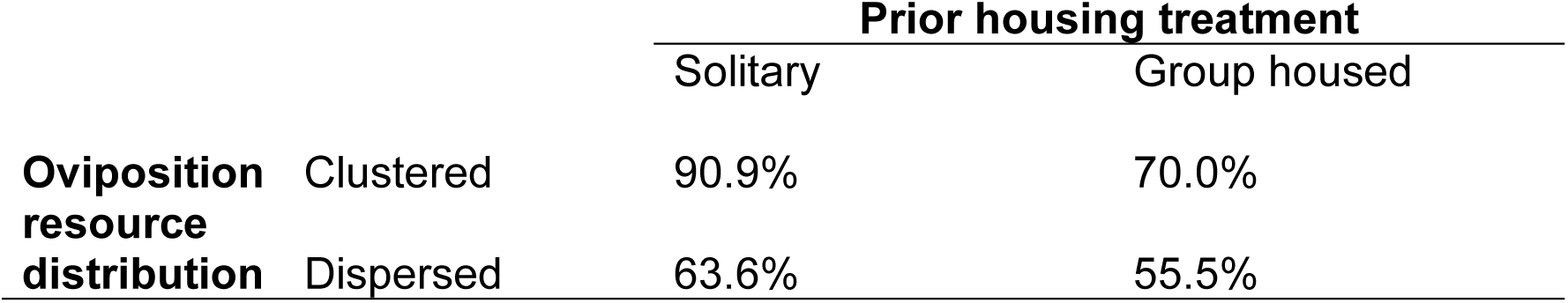
The effect of competition and resource distribution on the percentage of females who laid at least one egg

|  |  | **Prior housing treatment** |  |
| --- | --- | --- | --- |
|  |  | Solitary | Group housed |
| **Oviposition resource distribution** | Clustered | 90.9% | 70.0% |
|  | Dispersed | 63.6% | 55.5% |

Among those females that laid eggs, we found the opposite effect of these two treatments: prior housing significantly influenced clutch size, with group-housed females laying 22% fewer eggs than those from a solitary background (solitary: 18 ± 2 eggs; group housed: 14 ± 1 eggs; generalized linear model with quasipoisson errors F_1,69_ = 5.106, p = 0.027), but in this instance no significant effect of food patchiness was found (F_1,68_ = 0.073, p = 0.788). Again, there was no interaction between treatments (F_1,67_ = 0.076, p = 0.783) in this model.

Surprisingly, these effects of prior housing and egg-laying resource distribution did not lead to any difference between treatments in the number of adult offspring produced by the female (including all females that mated, whether or not they laid eggs: competition: F_1,99_ = 2.273, p = 0.135; oviposition substrate: F_1,98_ = 2.404, p = 0.124; treatment interaction: F_1,97_ = 0.064, p = 0.801). Neither treatment, nor the interaction, affected the sex ratio of offspring produced (F_1,85_ < 2.145, p > 0.147).

### Oviposition distribution and fitness amongst varying resource distributions

Females on dispersed resources spent less time on the food patches (34.2% of time on resources) than those on clustered resources (56.1%: raw number of records on food patches, linear model: F_1,99_ = 18.64, p << 0.001). Prior housing treatment had no equivalent effect (F_1,99_ = 0.321, p = 0.572), and there was no interaction between the two treatments (F_1,99_ = 0.155, p = 0.695).

There was no effect of housing treatment on the number of patches on which eggs were observed (χ^2^ test: χ^2^ = 5.44, p = 0.245), but females laid eggs on more of the available patches when these patches were clustered than when they were dispersed (χ^2^ = 32.06, p < 0.001; Fig. 6).

**Figure 6.**
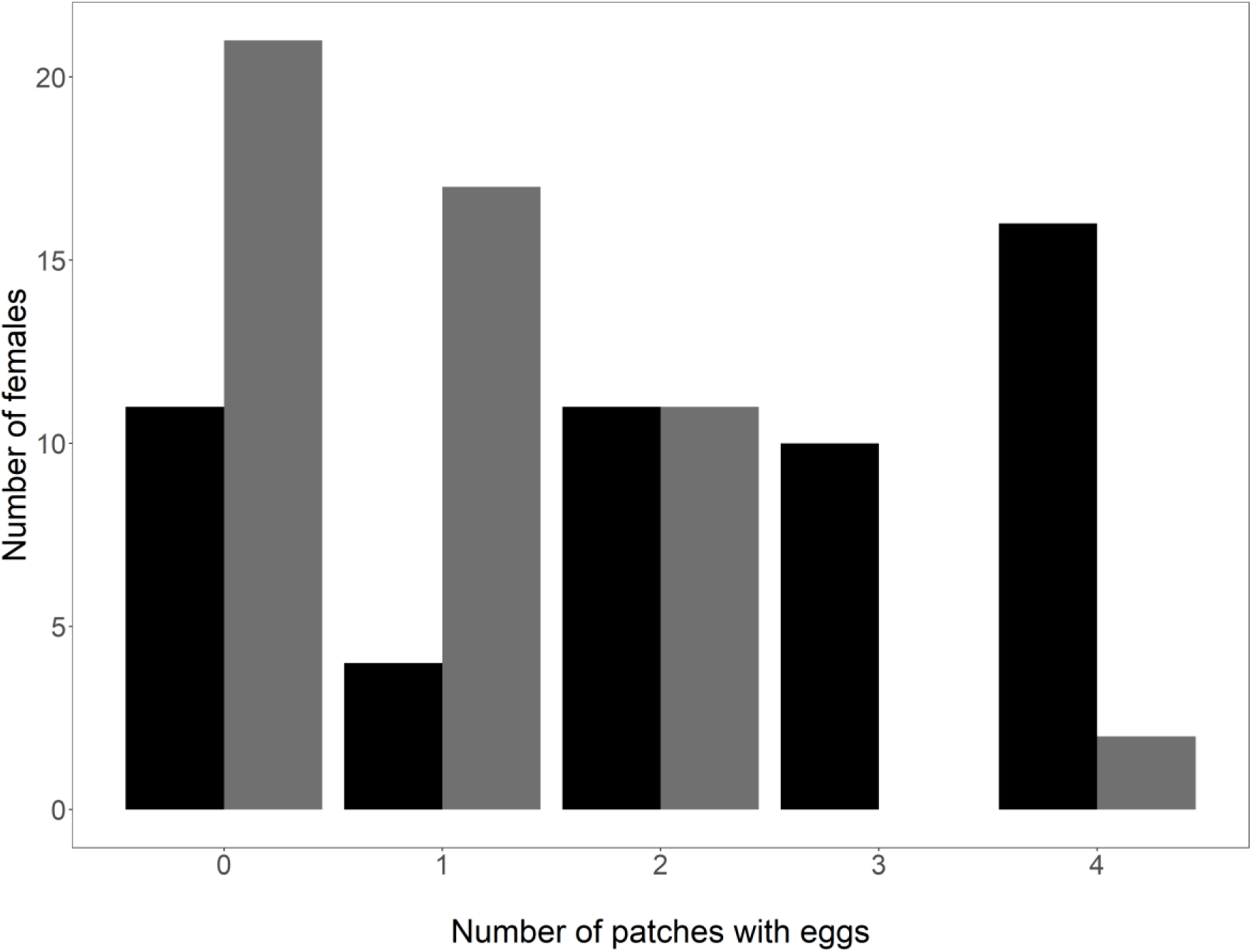
The number of females which laid on either zero, one, two, three or all four available oviposition patches in two different spatial arrangements. Black bars – clustered resources (N= 52 laying females); grey bars – dispersed resources (N = 51).

### Oviposition distribution on individual resource patches

Across all treatments, females were more likely to lay on the edge of patches than on the top surface (sum of eggs on sides vs top of 4 food patches; paired t-test, t = −6.426, p < 0.001); and this pattern held within each of the 4 treatment combinations (all p < 0.013).

### Post-treatment egg production

Of 101 females given the opportunity to lay in both Petri dishes and vials, 70 laid in both, 10 laid in neither, 20 laid none in the Petri dishes but did lay in the vials, and only 1 laid in the Petri dishes but none in the vials. The probability of laying in vials was thus significantly positively influenced by whether or not eggs had been laid in the Petri dish (GLM with binomial errors: χ^2^ = 20.8, p < 0.001), but not by either treatment (both p > 0.512). The number of offspring produced across both Petri dish and vial was not significantly affected by either prior housing density (χ^2^ = 18.39, p = 0.249) nor resource distribution in the Petri dish (χ^2^ = 20.47, p = 0.224).

## Discussion

### The impact of prior exposure to density on mating behaviours

Females that had been group housed before mating took longer to start copulating and copulated for a shorter duration than females maintained alone since emergence. Assuming both have at least an element of female control (Spieth, 1974, Mazzi et al., 2009), these behaviours suggest greater reluctance to mate among females from a group housed background. This matches findings from previous studies, which demonstrate that females are often more choosy in higher-density populations (Lehmann, 2007, Atwell and Wagner, 2014, Willis et al., 2011, Scott et al., 2020), where the risks of remaining unmated are lower and there is less pressure to mate with the first available male. Here, we interpret the delayed start of copulation and the shorter mating duration as indicators of choosiness – females from a grouped background showed lower willingness to mate with the first available male, and mated with him for less time, in expectation of future mating opportunities. While it is true that females in our grouped treatment encountered only other females, they may perceive this as evidence of generally higher population density, and hence greater likelihood of encountering multiple males.

It is notable that these density effects on females are the opposite to those demonstrated for males exposed to consexual rivals prior to mating, which stimulates more extended copulation durations in *D. melanogaster* and a number of other species (Bretman et al., 2009, Flay et al., 2009, García-González and Gomendio, 2004, Klemme and Firman, 2013). Male responses are interpreted as a reaction to a perceived increased risk of sperm competition, which males can best counteract by mating for longer – perhaps to increase the quantity of sperm transferred (Engqvist and Sauer, 2003, Simmons and Parker, 1992), but also possibly as a form of mate guarding (Vitta and Lorenzo, 2009). Because our measure of mating latency includes the time taken for males to initiate courtship, variation in mating latency might also be influenced by males’ reduced willingness to court females from group housed backgrounds. While there is evidence for females influencing copulation duration, it is clear that males can also affect this trait. So, an alternative explanation for the increased mating latency and reduced copulation duration we observed is that this variability is due to male rather than female behaviour. Males paired with previously group-housed females may have detected apparent high density of other females via pheromones remaining on the focal female: males are sensitive to the pheromones of other males carried on females (Friberg, 2006). While the responses of males to female density have not yet been explored in this species, the lengthened latency to copulate and subsequent shorted copulation duration could be due to males anticipating additional mating opportunities, influencing how much they invest in the focal female.

### Effects of prior housing density and resource spatial distribution on oviposition decisions and subsequent fitness effects

A higher proportion of females housed on clustered resources laid eggs, compared to those housed on dispersed oviposition resources. Those on clustered resources also laid on more of the available patches than those on dispersed resources. Clustered resources reduce the search time for females, and so they have more time available to lay eggs. This is supported by our data on the time spent on patches: females on the dispersed medium they spent less time on patches than females housed on clustered resources. However, the observation that some females did not lay at all, particularly on the dispersed patches, suggests that these may also be perceived as a less valuable resource than clustered. This may be arise from the fact that females apparently consider the degree to which larvae will need to travel when choosing oviposition sites (Schwartz et al., 2012): clustered patches better facilitate social aggregation in larvae, perhaps allowing for more efficient cooperative feeding (Dombrovski et al., 2017, Khodaei and Long, 2020). This difference was non-significantly exaggerated in the group housed females, which as noted above presumably have a higher expectation of future mating opportunities than the solitary females.

There may be physical environmental explanations for the difference in laying success on clustered vs dispersed resources. Like others, we found that females laid more eggs on the edge of the resource patches than on the top surface (Chess and Ringo, 1985, Moore, 1952). Although we did not control for the fact that patch side area is greater than patch top area, the available to lay upon are the same under both resource distributions. The number of sheltered edges is increased in clustered resources, and as *Drosophila* preferentially lay on the edges of resources, this increase in sheltered edges could be driving the preference for clustered egg-laying patches. Additionally, small patches of food are likely to dry out much more rapidly than larger patches, and clustering may help to mitigate this effect as well. A desiccated food resource will inevitably limit larval survival, giving a final explanation for females preferring this arrangement of patches. Further work will need to discriminate between these explanations for female behaviour.

While the distribution of egg-laying patches influenced the probability of eggs being laid and the number of patches laid on, this physical environment had no effect on the number of eggs laid. Instead, we found that prior housing treatment was important here: group housed females laid fewer eggs irrespective of egg-laying environment. Females engage in energetically expensive aggressive interactions with their consexual rivals in this species (Bath et al., 2017, Ueda and Kidokoro, 2002), perhaps leading to a trade-off in which group-housed females have less energy available for oviposition. In addition, when females oviposit in the presence of rivals, they copy their oviposition behaviours to reduce sampling costs (Malek and Long, 2020). In the absence of this information, females may have been slower to choose where to deposit their clutch. Finally, females from the group housed treatment are likely to anticipate a high level of competition for their offspring. This may have caused them to reduce the size of the clutch – perhaps increasing the size of eggs to improve their competitive advantage. However, although we did not measure egg size, we found no evidence to suggest there was a trade-off in the quality of eggs laid, or female investment per egg, as we found no treatment effects on the number of successfully eclosed adults. We recognize the possibility that fitness effects were absent because of the benign laboratory conditions (e.g. *ad libitum* food and constant temperature), and future work should test for fitness effects in a more stressful environment and over subsequent generations.

It is also possible that fitness effects were not detected because our experimental design required the removal of the females before they had completed oviposition of all fertilised eggs: this was necessary so that the location of oviposited eggs could be recorded, and larvae hatch after 22-24 hours at ∼25°C (Fernández-Moreno et al., 2007, Markow et al., 2009). But if females were given the opportunity to continue ovipositing on the patchy resources, it is quite plausible that their choice of egg location would have impacted overall fitness.

We have demonstrated that female density has significant effects on mating and egg-laying behaviour. Females from group-housed conditions are slower to accept mating and mate for less time, which we suspect is related to future opportunities to mate, and lay fewer eggs, which is likely to be due to anticipated competition for their larvae. While the physical arrangement of egg-laying patches also has significant effects on oviposition behaviour, with clustered resources being preferred over dispersed ones, there is no interaction between this physical stimulus and the social stimulus of perceived population density. While reproduction-related effects of density are well known in males, equivalent study in females has so far been lacking. We have demonstrated that social environment can also have profound effects on females too: how the social environmental affects the sexes simultaneously may be of particular interest in the future.

## Acknowledgments

EC, JB, and MT conceived the study with advice from CD, EC conducted the experiments, EC and MT analysed the data, and all authors contributed to writing the manuscript. Thanks to Fliss Thom for help with fly and laboratory maintenance and experimental support.

